# Attentional guidance through object associations in visual cortex

**DOI:** 10.1101/2024.02.02.578555

**Authors:** Maëlle Lerebourg, Floris P. de Lange, Marius V. Peelen

## Abstract

Efficient behavior requires the rapid attentional selection of task-relevant objects. Preparatory activity of target-selective neurons in visual cortex is thought to support attentional selection, guiding spatial attention and favoring processing of target-matching input. However, naturalistic searches are often guided by non-targets, including target-associated “anchor” objects. For instance, when looking for a pen, we may direct our attention to the office desk on which we expect to find it. Here, using fMRI and eyetracking in a context-guided search task, we tested whether preparatory activity in visual cortex reflected the target, the guiding anchor object, or both. Participants learned associations between targets and anchors, reversing across two scene contexts, before searching for these targets. Participants’ first fixations were reliably guided by the associated anchor. Preparatory activity in lateral occipital cortex (LOC), and right intraparietal sulcus (IPS), represented the target-associated anchor rather than the target. These results shed light on the neural basis of context-guided search in structured environments.

## Introduction

Imagine yourself standing in your colleague’s messy office, looking for a pen to write down a note. Visual search tasks like these are challenging, yet we perform them frequently, efficiently, and seemingly without much effort. A seminal finding in the field of visual search is the finding that attentional selection of a target object is supported by preparatory activity of target feature-selective neurons in visual cortex (Desimone & Duncan, 1995), with preparatory activity targeting higher-level object representations in object-selective regions of the ventral stream when searching for objects in naturalistic scenes (Battistoni et al., 2017). For example, when monkeys were cued to search for a specific object, the firing rate of neurons in monkey inferotemporal cortex (IT) tuned to this object increased during the interval between the cue and the search display (Chelazzi et al., 1993a, 1998). Similarly, in humans, fMRI activity patterns in visual cortex, especially object-selective regions within the ventral visual stream, represented the cued target in the absence of visual stimulation. This preparatory activity has been interpreted as an “attentional template” – a top-down bias in favor of target-matching input, resolving the competition between multiple visual stimuli, guiding attention and eye movements, and enhancing processing of potential targets (Bichot et al., 2005; Desimone, 1998; Eimer, 2014; Wolfe, 1994).

However, search in daily life differs from search in previously employed laboratory tasks in important ways. For example, in structured real-world scenes, search is not only guided by target features but also by contextually associated (non-target) objects (Boettcher et al., 2018a; Koehler & Eckstein, 2017a, 2017b; Mack & Eckstein, 2011; Zhou & Geng, 2024). Going back to our example of looking for a pen, we may direct our attention to the office desk on which we expect the pen to be. In this case, the office desk reflects a so-called ‘anchor’ object (Boettcher et al., 2018b; Draschkow & Võ, 2017; Helbing et al., 2022): it is large, salient (i.e., easy to find), associated with the pen, and provides spatial predictions about the pen’s location. Whether or how preparatory activity can support context-guided search has not been previously investigated. Recent theories of visual search (Wolfe, 2021; Yu et al., 2023) distinguish between ‘guiding templates’ and ‘target templates’, but their respective link to preparatory activity is not yet clear (Dodwell & Eimer, 2024). We reasoned that if preparatory activity in visual cortex serves as an attentional guidance mechanism, it should not necessarily reflect the target, but rather those features or objects that are effective for guidance. If, instead, preparatory activity primarily supports target identification and/or processes supporting target-related decision making, it should reflect features of the target.

To test this, we designed a context-guided search task that allows for separating preparatory fMRI activity related to the target and to the anchor. Participants in the fMRI experiment were first familiarized with novel target-anchor associations, learning to associate two target categories (books and bowls) with two different tables across two rooms (e.g., the book was found on Table 1 in Room 1 but on Table 2 in Room 2; Figure 1). This design allowed us to test whether relevant guiding objects in the current context were represented in preparatory activity. The tables in our scenes were useful for search in a similar way as real-world anchors are, being easy to find and providing spatial predictions constraining the search space for the target, while controlling for visual or semantic similarity. After familiarization, participants searched for the peripherally presented target objects. To measure preparatory activity for the target and anchor objects, we introduced preview-only trials, in which participants prepared to search for a target object within a given room, but the target and anchor objects did not appear in that trial. These trials allowed us to isolate fMRI activity patterns related to search preparation, and test whether those carried information about the associated anchor or, alternatively, only the current target.

**Figure 1:**
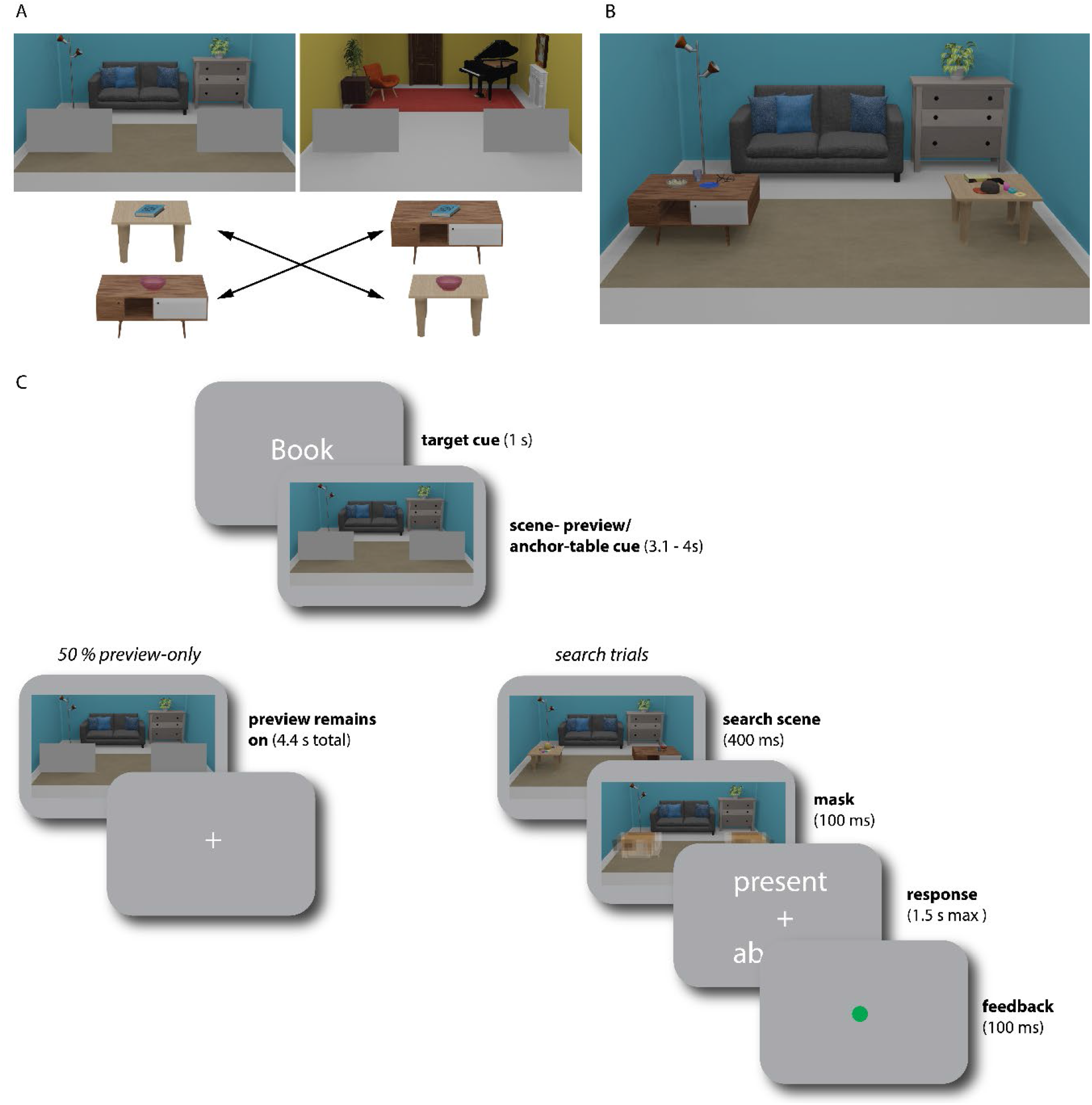
Experimental design and timeline. **A** Participants learned room-dependent associations between target objects (books and bowls) and two different tables serving as anchor objects. **B** Example of a search scene. Both tables could either appear left or right in the scene, with multiple objects placed on them. If the associated target was present, it would appear on the associated table. Across the two rooms, associations between targets and anchors were switched. **C** Timeline of a trial. On each trial, participants were cued to search for one target category, followed by a scene preview. In search trials, a search scene would then briefly appear and participants were free to move their eyes. At the end of the trial, they indicated whether the cued target had been present. In half of the trials, however, participants prepared to search, but no search scene appeared after the preview and no response was required.

Following previous work (Gayet & Peelen, 2022; Peelen & Kastner, 2011) our analyses focused on two visual cortex regions that may encode preparatory attentional templates: object-selective lateral occipital cortex (LOC) in the ventral visual stream and early visual cortex (EVC). Specifically LOC has most consistently been implicated in visual search for real-world targets (Gayet & Peelen, 2022; Peelen & Kastner, 2011; Soon et al., 2013; van Loon et al., 2018) and may therefore also represent a guiding template for anchor objects.

We found that the anchors guided eye movements, with the first fixation in the scene already directed towards the target-associated anchor, even in target-absent trials. Crucially, we found that preparatory fMRI activity in LOC during search preparation (preview-only trials), represented the target-associated anchor, independent of the target object and independent of the room that participants were searching. These results demonstrate that preparatory activity in visual cortex supports context-guided search by reflecting features relevant for guidance.

## Results

Participants (N=34) searched for target objects (books and bowls) within two scene contexts (3D rendered images of a blue and yellow living room, see Figure 1) while undergoing fMRI scanning. The target objects were small and appeared in the periphery, placed among other non-target objects on one of two tables in the room. Within each room, targets of a given category always appeared on the same table, and the tables therefore acted as anchor-objects for the targets (100% validity). Across rooms, the target-anchor associations were switched (e.g., blue living room: book on Table 1, bowl on Table 2; yellow living room: book on Table 2, bowl on Table 1; specific associations counterbalanced across participants). Participants were familiarized with these associations before fMRI scanning (see Anchor-target association training). On each trial, a word cue (‘book’ or ‘bowl’) indicated the upcoming search target (1s), followed by a preview of the blue or yellow living room (3.1 – 4s), in which the tables were still occluded by grey rectangles. Together with the target cue, this preview could be used to prepare for the specific anchor-table associated with the target in the current room. Critically, within each room, the same table could appear either left or right with equal probability, and the preview was therefore not spatially predictive. All trial types (e.g., trials with different targets and rooms) were randomly intermixed, therefore requiring participants to retrieve the relevant target-anchor associations anew on each trial. On half of the trials (*search trials*), the occluders were then removed, briefly revealing the anchor-tables and objects placed on them. On all search trials, four different objects were placed on each table. On target absent trials, the target was substituted by an additional distractor object (e.g., a laptop, a coffee mug, or playing cards).

The search target was present on 50% of those trials. During this search phase, participants could freely inspect the scene and move their eyes. Average accuracy on those search trials was 72.46% (see Supplementary Materials for behavioral analyses). This shows that the search task was indeed challenging, making a preparatory template and contextual information useful for the task. Afterwards, a response screen appeared, prompting participants to report whether the target had been present or absent. On the other half of trials (*preview-only trials*), however, the occluders remained, and thus no objects or associated tables were shown. On those trials, participants were asked to keep fixating at the image center and no response was required. Importantly however, since it was unknown during the preview whether the search scene would appear or not, participants still had to prepare to search on every trial. The preview-only trials were the trials of interest for the fMRI decoding, allowing us to isolate preparatory fMRI activity.

## Eyetracking analysis

### First fixations are guided by anchor objects

To test whether anchor objects guided attention, we analyzed eye movements during the search phase, focusing on the first fixation in the scene (mean onset: 229.53 ms (sd 29.20)) as an index of overt attentional guidance. Target absent trials, in which target features could not guide fixations provided a pure measure of guidance by the anchor. On those trials, 19.20% (sd 19.44) more first fixations were directed towards the correct anchor compared to the other non-associated anchor (CI = [12.50, 24.70], p < 0.001; Figure 2A). On target present trials, 25.24% (sd 20.59) more fixations were directed towards the correct anchor than the other non-associated anchor (CI = [19.00, 32.47], p < 0.001; Figure 2A), which was significantly higher than for target absent trials (CI = [0.40, 11.41], p = 0.03). This indicates that anchor features indeed guided attention, with additional guidance provided by target features when they were present. Finally, guidance by anchors also yielded search benefits, as indicated by a significantly positive correlation between anchor guidance and d’: participants with more selective (anchor-guided) first eye movements on target absent trials also had a higher d’ (r = 0.55, p < 0.001; Figure 2B). This correlation was also significant for target present trials (r= 0.60, p < 0.001; Figure 2B).

**Figure 2:**
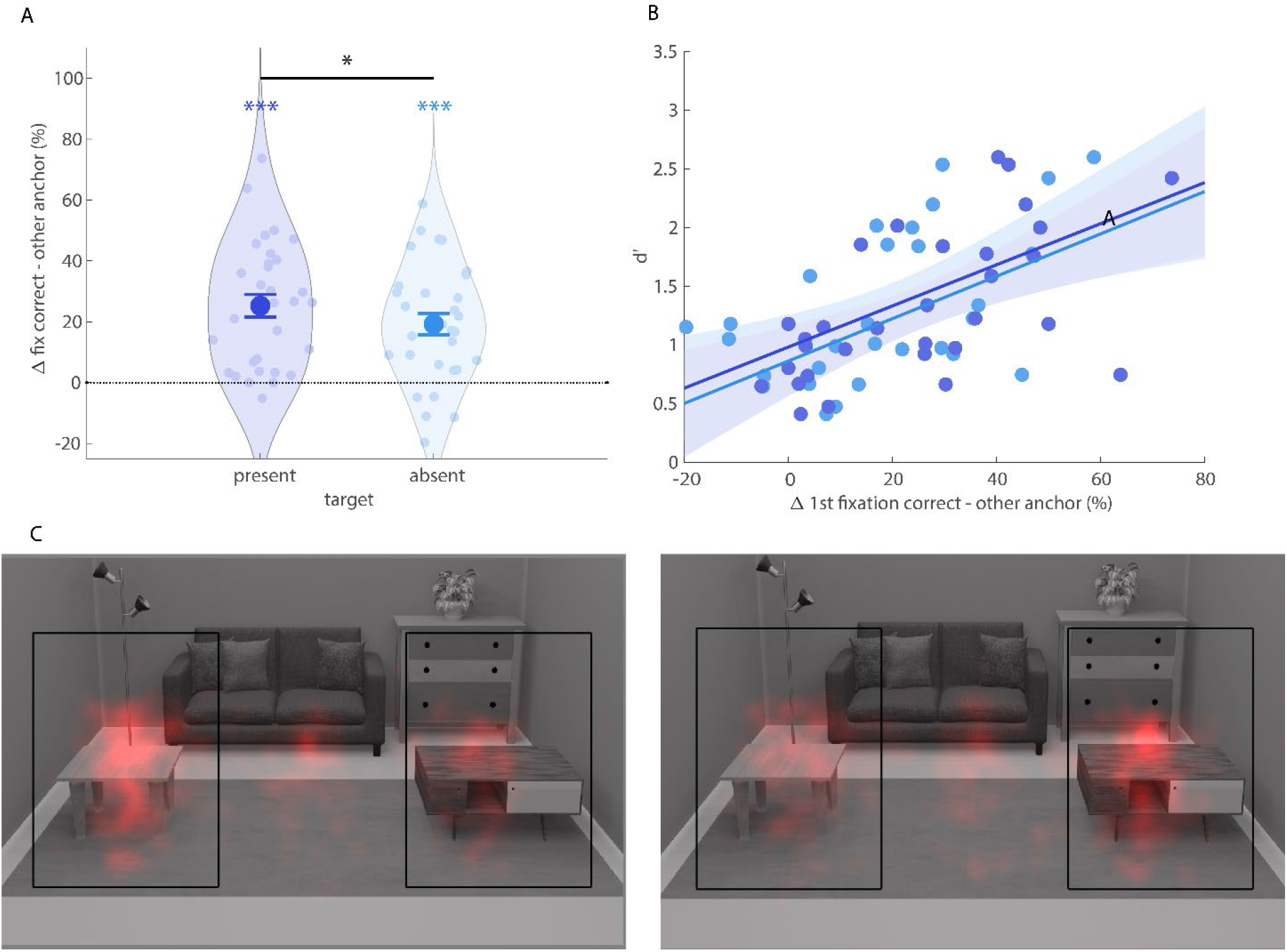
Eye tracking results. **A** Difference in percentage of first fixations directed towards the correct (relevant) anchor and the other (irrelevant) anchor for target present trials (left) and target absent trials (right). All error bars are SEM. ***, p<.001 **B** Correlations of d’ during search with first fixation differences on target present (dark blue) and target absent (light blue) trials across participants. Shaded areas shows 95% confidence intervals. **C** Heatmaps for the first fixations on target absent trials when the associated anchor was on the left (left panel) or right (right panel). The black squares indicate the two areas of interest (AOIs) used for eyetracking analysis and were not visible for participants.

## fMRI results

### Preparatory activity in LOC does not reflect the target

Having established reliable guidance by anchors, we turned to investigate preparatory fMRI activity patterns on the preview-only trials within two regions of interest: EVC and LOC. First, we tested whether there was any information about the target object participants prepared to search for (Figure 3A). We trained classifiers to decode the target category (book or bowl) of each trial, collapsed across scene contexts and therefore also across associated anchors. For both ROIs, we intersected group-level masks with voxels sensitive to target and/or anchor features at their retinotopic locations in the task. To ensure the robustness of our results to different voxel selections, we repeated this analyses across ROIs of different sizes (see Figure 3). Classifier performance was calculated based on the distance from the decision boundary, providing a more sensitive and continuous estimate compared to classification accuracy based on binary labels (Walther et al., 2016). This was calculated as follows: 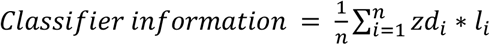. For each beta *i*, the z-scored distance-to-bound (*zd*_*i*_) was multiplied by its correct label (*l*_*i*_ ∈ [−1, 1]) and this score averaged across all *n* betas (see Ref. 20 for a similar approach). All our main results were replicated with accuracy-based analyses (see Figure S3.).

**Figure 3:**
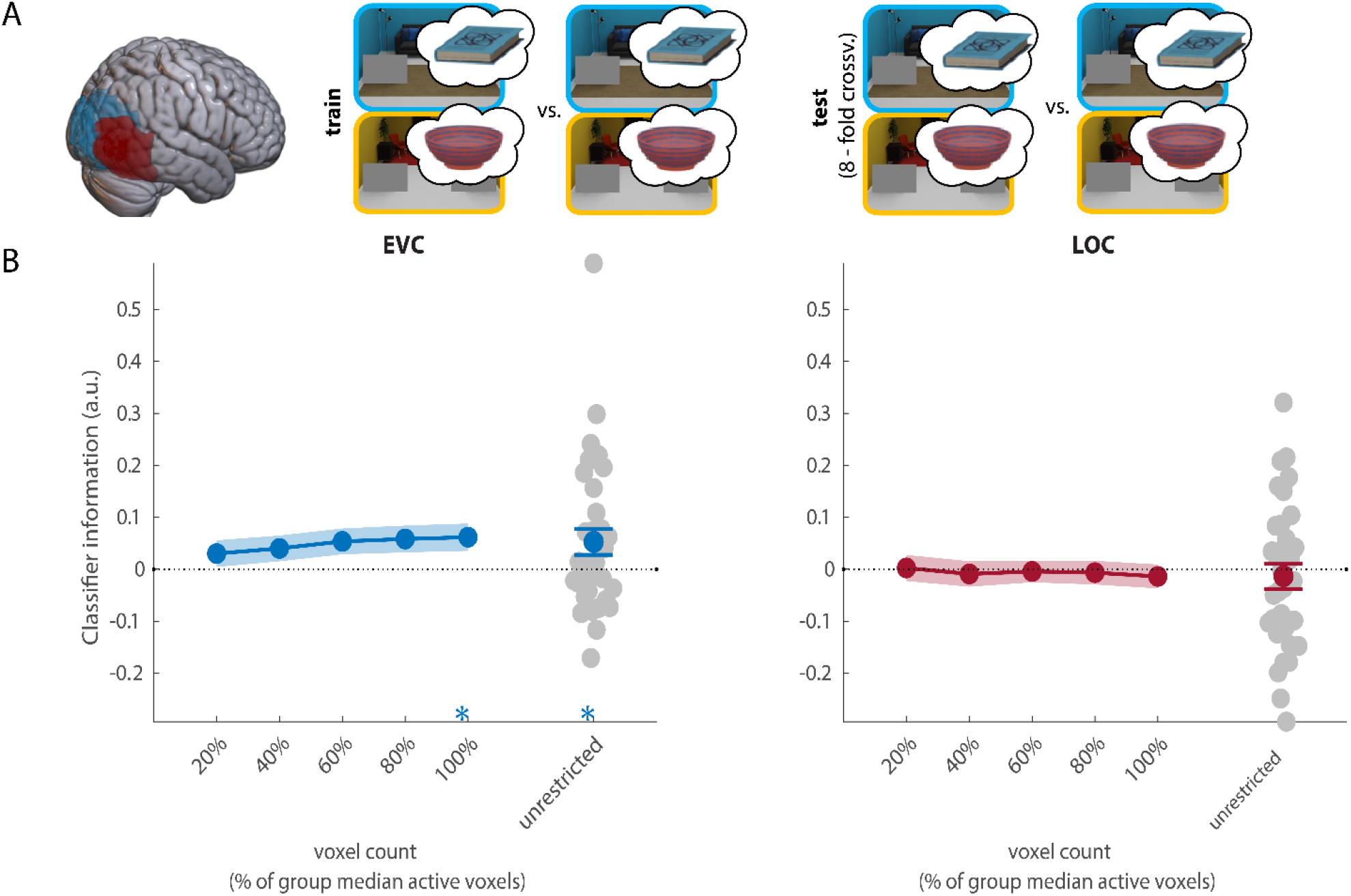
Decoding of the target object within search task runs. **A** Overview of the decoding scheme, decoding the target (book/bowl) from the preview only trials (leave one run out cross-validation). Brain shows anatomical location of the ROIs. **B** Decoding results in EVC and LOC for all selective voxels of individual participants (unrestricted -ROI) and sub-ROIs of different sizes. Grey dots show decoding for individual participants in the largest (unrestricted) ROI. All error bars are SEM. *, p<.05. For sub-ROIs, asterisks show significance after threshold-free cluster enhancement (TFCE).

Within EVC, the current search target could be decoded above chance (p = 0.02, CI = [0.0078, 0.11]; Figure 3B), but this was not consistent across a majority of voxel selections and should therefore be interpreted with caution. Notably, in contrast to previous studies, there was no information about the current search target in LOC (p = 0.58, CI = [-0.06, 0.04]). Taken together, these results indicate that the search target was not strongly represented in preparatory activity during context-guided search. This result is in line with the ineffective attentional guidance offered by target features in this task. As attention (and eye movements) were primarily guided by anchor objects, we expected that preparatory activity would represent these anchor objects.

### Preparatory activity for associated anchor objects

Next, we turned to our main analysis, testing whether preparatory activity indeed reflected the target-associated anchor that guided eye movements. For this analysis, classifiers were trained to decode the target-associated table (table 1 vs table 2) from fMRI activity patterns in preview-only trials (see Figure 4A). There was no evidence for anchor-information in EVC (p = 0.24, CI = [- 0.03, 0.14]; Figure 4B). Importantly, however, the associated anchor could be reliably decoded from LOC (p = 0.001, CI = [0.03, 0.15]; Figure 4B). Decoding was consistent across all ROI sizes (Figure 4B). Note that there was no external visual cue for the relevant table and this preparatory activity must therefore have been internally generated, based solely on the learned target-anchor associations.

**Figure 4:**
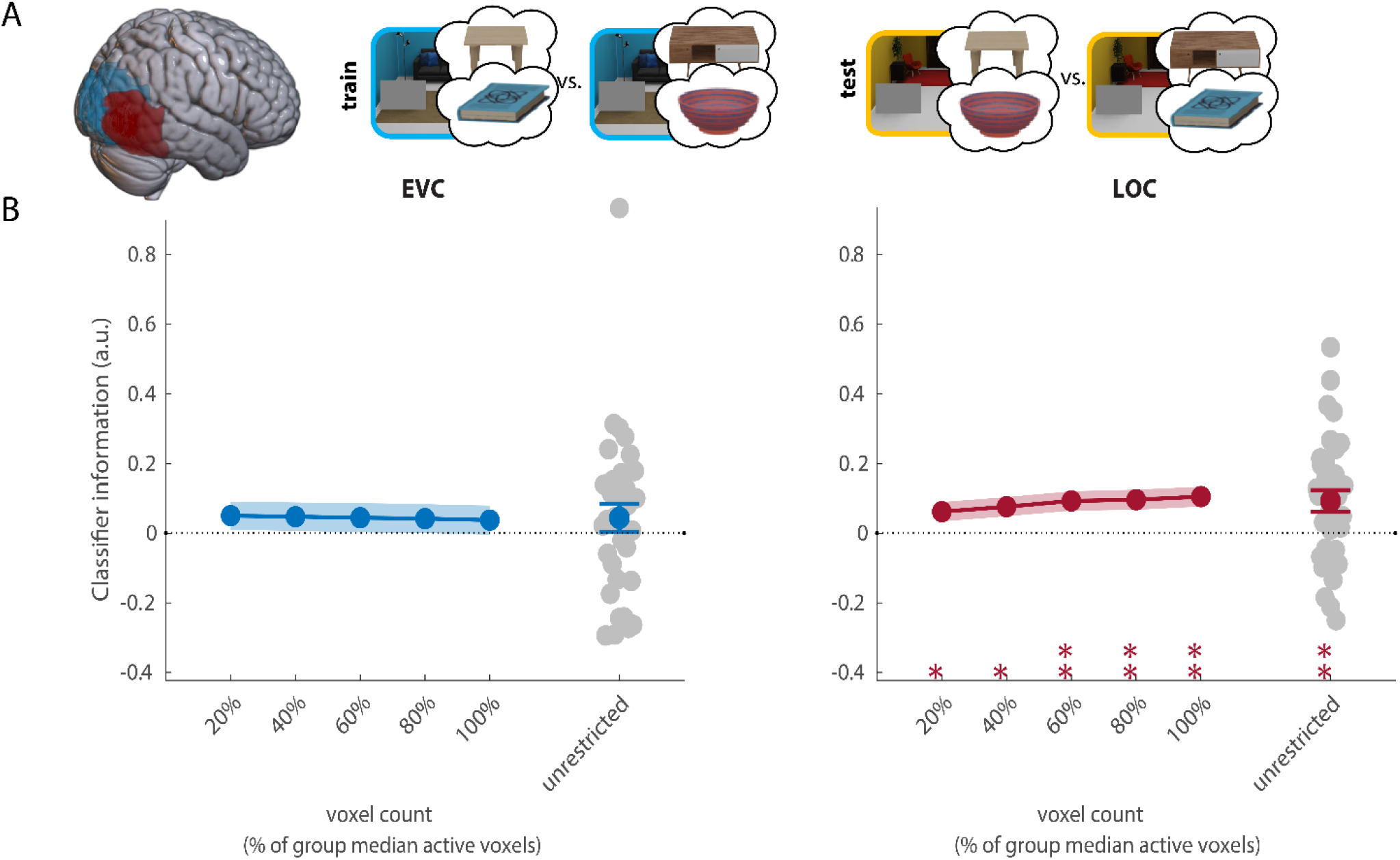
Decoding of the associated anchor within search task runs. **A** Overview of the decoding scheme, decoding the associated anchor (Table 1/Table 2) from the preview only trials (leave one run out cross-validation). Brain shows anatomical location of the ROIs. **B** Decoding results in EVC and LOC for all selective voxels of individual participants (unrestricted-ROI) and sub-ROIs of different sizes. Grey dots show decoding for individual participants in the largest (unrestricted) ROI. All error bars are SEM. **, p<0.01; ***, p<.001. For sub-ROIs, asterisks show significance after threshold-free cluster enhancement (TFCE).

### Cross-scene decoding analysis: Anchor templates in LOC generalize across contexts

Since the associated anchor could be reliably decoded during search preparation, we next asked whether preparatory anchor activity was independent of the cued target and the specific room (Figure 5A). Thus, we trained classifiers on dissociating the currently relevant anchor in one room, testing whether they could also decode the relevant anchor in the other room, where the target-anchor associations were reversed. This analysis additionally allowed us to rule out that differences in anticipated search difficulty between anchors contributed to anchor-decoding, as the performance differences we observed were scene-specific (see Behavioral analyses in Supplementary Materials).

**Figure 5:**
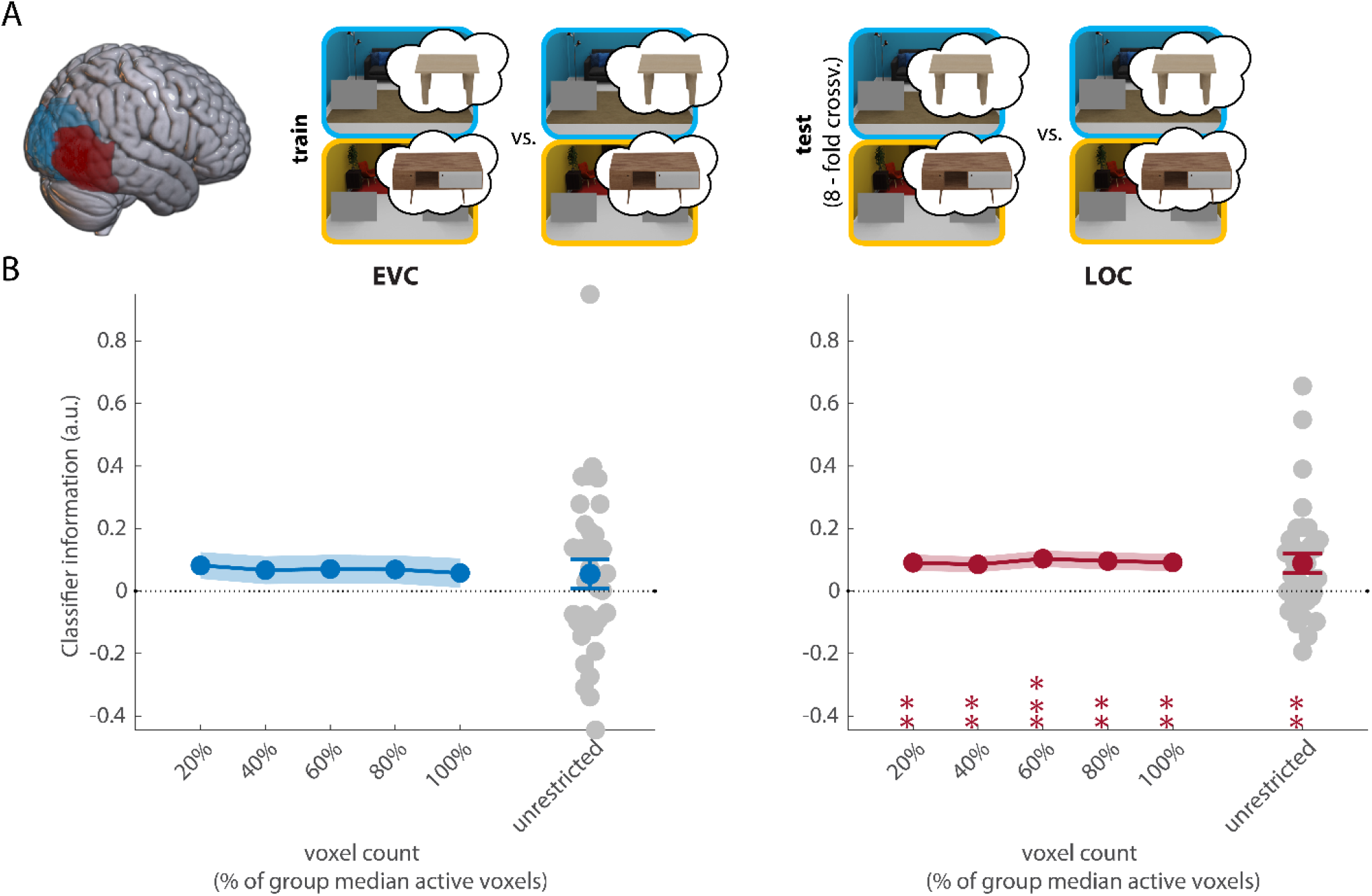
Decoding of the target-associated anchors across rooms. **A** Overview of the decoding scheme, decoding the associated anchor (table 1/table 2) from the preview only trials in one room/scene context, testing on the other (cross-scene decoding). Brain shows the anatomical location of the ROIs. **B** Decoding results in EVC and LOC for all selective voxels of individual participants (unrestricted-ROI) and sub-ROIs of different sizes. Grey dots show decoding for individual participants in the largest (unrestricted) ROI. All error bars are SEM. *, p<0.05; **, p<0.01. For sub-ROIs, asterisks show significance after threshold-free cluster enhancement (TFCE).

In this analysis, above-chance decoding would indicate preparatory activity for the anchor table, generalizing across associated targets, while below chance decoding would indicate preparatory activity for the target (independent of the associated anchor table). Within EVC, preparatory activity did not generalize across rooms (p = 0.28, CI = [-0.031, 0.13]; Figure 5B). Importantly however, the anchor template in LOC was independent of scene context and the associated target (p = 0.004; CI = [0.03 0.15]; Figure 5B). Decoding was consistent across ROI sizes (Figure 5B).

Overall, this result shows that preparatory activity in LOC specifically reflected the anchor, irrespective of the associated target and independent of the room in which participants prepared to search.

### Searchlight analyses

To complement the ROI analyses, we conducted a searchlight analysis to test whether additional brain regions showed context- and target-independent preparatory activity reflecting the anchor, as we found for LOC. This analysis revealed one cluster with above-chance decoding, located within the right intraparietal sulcus (IPS; Figure 6). We did not observe significant clusters within LOC, likely explained by reduced sensitivity of this searchlight analysis to spatially distributed activity patterns, which may have additionally varied across participants, together with a stringent multiple comparison correction (threshold-free cluster enhancement (TFCE; Smith & Nichols, 2009)). Indeed, a whole-brain searchlight analysis at a more lenient threshold (p < 0.005, uncorrected) revealed clusters of voxels within LOC (Figure S4). There were no significant clusters showing negative cross-scene decoding (which would be indicative of preparatory target activity).

**Figure 6:**
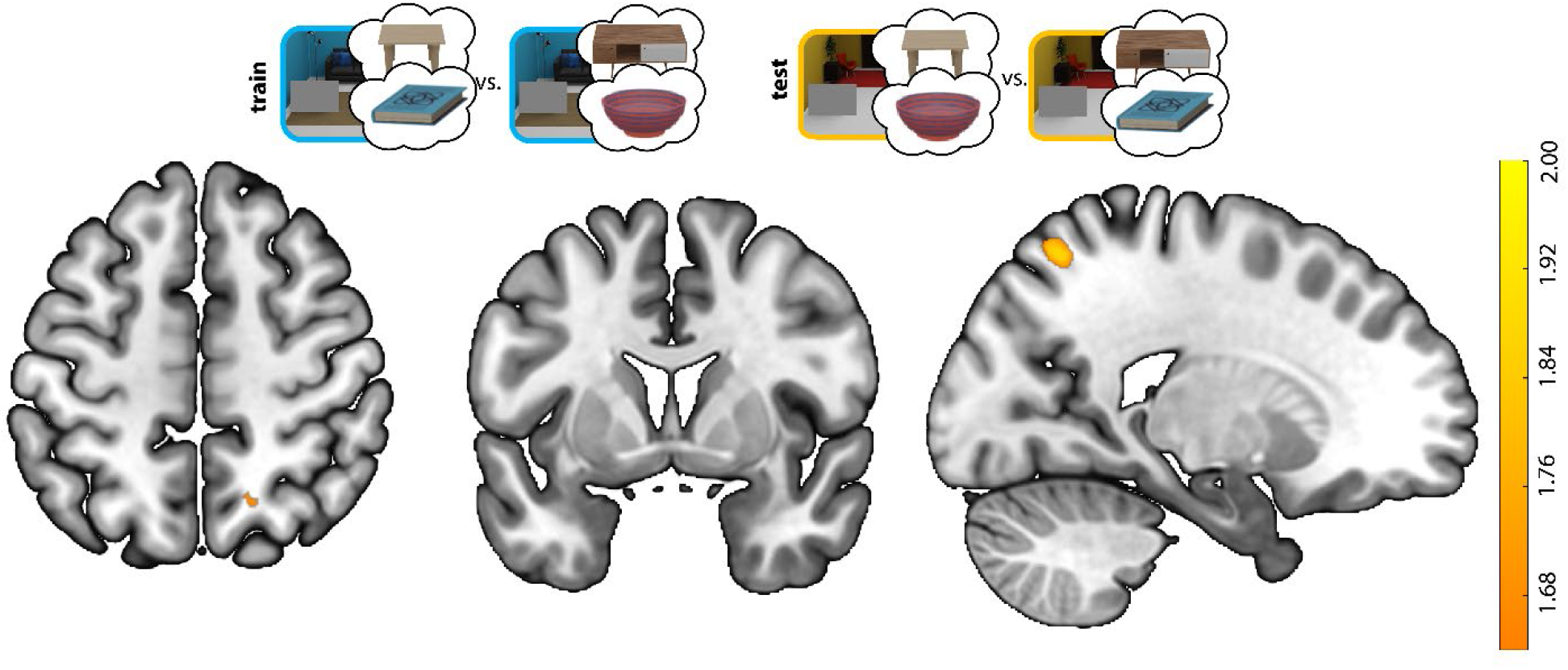
Searchlight for decoding the target-associated anchor across scene-contexts. The searchlight analysis revealed a cluster in the right intraparietal sulcus (x, y, z = 20.1, -63.6, 58, 203 mm^3^). Color indicates z-scores after threshold free cluster enhancement (TFCE).

## Discussion

Attention in structured, real-world scenes is often guided by features that are not based on a precise, veridical representation of the target (Wolfe, 2021; Yu et al., 2023), and can even be based on entirely different non-target objects, as in the case of guidance by anchor objects. Here, we tested whether and how preparatory activity supports such context-guided search.

Our results show that preparatory activity patterns in LOC, previously implicated in encoding preparatory attentional templates for naturalistic search targets (Gayet & Peelen, 2022; Peelen & Kastner, 2011), reflects relevant guiding objects in the current scene context, rather than the search target per se. Importantly, in contrast to the target object in this or previous studies investigating preparatory activity, the guiding object did not have to be reported, nor was it explicitly cued. Our findings are in line with recent visual search theories (Wolfe, 2021; Yu et al., 2023) that propose a dissociation between templates for attentional guidance and templates for later target processing, where a veridical and precise target representation is mostly useful for those later decision stages. By disentangling target and guiding features, our results indicate that preparatory activity in LOC supports guidance itself. Besides target report, the current design also allowed us to dissociate this biasing signal from the external target cue. Such cues are seldomly encountered in real-world searches and the representations based on external cues (even when they are abstract) can differ from internally generated ones, e.g., activate more posterior, purely visual regions (Hu & Yu, 2023). In the current study, the same target cue was associated with different anchors depending on the room, such that the associated anchor had to be retrieved from memory based on learned associations. Overall, our results show that preparatory activity in visual cortex reflects an internally generated guiding template, independent of external cues or target report.

We found that preparatory activity in visual cortex can be highly flexible and context-dependent, representing a different anchor when looking for the same target in different scene contexts (blue or yellow living room). Across rooms, each target was equally associated with either table; which anchor was effective for guidance depended on the combination of target and scene context. Furthermore, all trials were randomly intermixed in the experiment. The target-anchor associations therefore did not reflect general semantic associations, but rather context-specific associations for guidance, also independent of priming and any visual similarity between targets and anchors. Despite this reversal across contexts, associations were quickly learned, used for eye movement guidance, and reflected in preparatory activity, thereby demonstrating the flexible top-down nature of preparatory activity. This flexibility would be beneficial for search in daily life, where objects can appear next to different anchors depending on the context (e.g., a coffee cup next to the coffee machine in a kitchen but next to the monitor in an office).

What is the source of those flexible and context-dependent biases in visual cortex? Previous research investigating learned associations between scene context and target location (e.g., learning that target 1 usually appears in the top left corner of scene 1) has shown that guiding attention based on memory is mediated by interactions between hippocampus and visual cortex (Summerfield et al., 2006). These interactions have been shown to influence visual cortex activity prior to stimulus onset (Favila & Aly, 2023; M. G. Stokes et al., 2012; Summerfield et al., 2006). Similar mechanisms may be involved when recently learned context-dependent object associations are retrieved for guidance, as in the current study. It is possible that a different mechanism supports guidance for highly stable object-associations that have been learned over many years (e.g., toilet roll holder next to toilet) as such associations may affect the cortical representation of the associated objects, leading to a more integrated representation of object pairs within object selective visual cortex (Kaiser et al., 2019).

The searchlight analysis revealed that anchor-specific preparatory activity was also present in the right intraparietal sulcus. Similar to LOC, this activity generalized across target identity and scene contexts. IPS may be well suited to encode such guiding templates, as neurons in this region are shape sensitive (Konen & Kastner, 2008) and are involved in working memory (Christophel et al., 2015, 2018; Iamshchinina et al., 2021; Rademaker et al., 2019) and eye movement control (Grefkes & Fink, 2005). IPS neurons have also been implicated in “information sampling” (Gottlieb et al., 2013; Gottlieb & Oudeyer, 2018), encoding the expected information gain of attending to a particular object (Foley et al., 2017; Vossel et al., 2006, 2015). Within naturalistic scenes, both IPS and LOC have been found to encode expected spatial locations of targets, thereby supporting contextual guidance in scenes (Preston et al., 2013). This could suggest that these regions are sensitive to both the location and the identity of guiding objects. Future research targeting IPS and its subregions more specifically is needed to confirm these findings.

While we found no evidence for preparatory activity related to the target object in LOC, participants must have correctly prepared for the cued target, as they performed above chance and target features improved guidance. It is possible that such activity starts later, once the associated anchor has been located and participants need to decide whether the search target was present or absent. In the current experiment, this stage could not be separately investigated. In line with this, there is evidence from sequential search and working memory tasks that items relevant in the future are not (or less well) decodable in visual cortex, compared to immediately relevant items (Christophel et al., 2018; e.g., van Loon et al., 2018). Future studies could use our paradigm in combination with time-sensitive methods (e.g., EEG/MEG) to test for sequential anchor-target activity. Alternatively, the current paradigm could be adjusted for fMRI to measure preparatory activity after the anchor has been located, for example using a two-step preview task. More generally, while other, more sensitive designs or analyses might reveal concurrent target and anchor templates, this would not change our main conclusion that preparatory activity can represent non-target features based on learned associations.

Our results support the view that attentional templates, especially guiding templates, are flexible and represent useful guiding information rather than necessarily a veridical and detailed representation of the target (Yu et al., 2023). Whether a guiding template reflects target or non-target features is most likely a trade-off, depending on the relative ease with which target features themselves can be effectively used for guidance (influenced by, e.g., saliency, predictability, target-distractor similarity), and the degree to which non-target features are associated with the target (making them more or less informative). In this study, our aim was to test whether preparatory activity reflected such non-target features in situations where these provided an advantage for guidance. If guidance by target features was relatively more effective, e.g., because target features were more salient or target-anchor associations were unreliable, it is likely that preparatory activity would more strongly reflect the target, as reported previously (Chelazzi et al., 1993b; Gayet & Peelen, 2022; Peelen & Kastner, 2011; M. Stokes et al., 2009). Testing this trade-off in detail remains a relevant question for future research, as object associations in the real world vary in strength across target-anchor pairs and are not always deterministic. At the same time, it is likely that anchor templates in real-world search are relatively reliably activated during search because of our extensive experience with real-world object associations (Kallmayer et al., 2024).

The current study allowed us to isolate one key aspect of naturalistic search - guidance by target-associated objects in the scene - and test the content of preparatory activity using a controlled design. The tables in our task were similar to real-world anchors in terms of being large and salient objects that provided spatial predictions about target locations, thereby constraining search. However, our study does not capture all aspects of real-world anchor search. For example, real-world target-anchor associations are typically learned over many years (e.g., toothbrush-sink) and are characterized by high semantic, functional, and/or contextual similarity of the objects involved, which are difficult to tease apart in an experiment. Besides guidance by anchor objects, naturalistic scenes also contain a wealth of other regularities that can be quickly extracted and support search (Hollingworth, 2012; Peelen & Kastner, 2014; Võ, 2021; Wolfe et al., 2011). For instance, scenes provide contextual expectations about target appearance that can shape the preparatory template in visual cortex (Gayet & Peelen, 2022). Target location may also be predicted based on the overall spatial layout of the scene, guiding attention and eye movements (Castelhano & Heaven, 2011; Neider & Zelinsky, 2006). Furthermore, frequently co-occurring objects may be grouped together, reducing competition between objects in the scene when those objects act as distractors (Kaiser et al., 2014). Rather than fully recreating all aspects of real-world search, the current findings reflect a proof-of principle of a key aspect of context-guided search, showing how preparatory activity can support context guidance based on learned object-associations.

In conclusion, we show that context-guided search is supported by self-generated and context-dependent preparatory biases in object-selective cortex, clarifying the functional role of preparatory activity during visual search. The ability to quickly learn novel object associations and flexibly adapt preparatory biases to represent the most useful features for attentional guidance is an important factor contributing to the efficiency with which we select and interact with relevant objects in our daily life environments.

## Materials and Methods

### Participants

Thirty-four participants (22 women, mean age: 23.5, sd: 5.8) from the Radboud University subject pool were included in this study. Seven additional participants were tested, but not included in the final data set: 5 due to at-chance accuracy in the search task (determined by a 1-sided binomial test with α = 0.05), one chose to abort the experiment and one due to excessive head movement. Those participants were replaced until reaching the planned sample size of 34, sufficient to uncover an effect of medium size (*d* = 0.5) with 80% power. All participants received 25€ for their participation and provided written informed consent, declared themselves free of epilepsy, and had normal or corrected-to-normal vision. All procedures were approved by the local ethics committee (CMO region Arnhem-Nijmegen, the Netherlands, Protocol CMO2014/288).

### Search task

Participants completed 256 trials of the task in the fMRI scanner (8 runs of 32 trials each). They searched for two different types of targets (books or bowls) in two 3D rendered living room scenes (see Figure 1).

Each trial began with a target cue (‘Book’ or ‘Bowl’) written on screen (1s), followed by a living room preview, with grey occluder rectangles placed at the location of the tables, slightly below fixation and in the left and right periphery. During target cue and scene preview, participants were asked to keep their gaze on a central fixation cross. The preview remained on for 4.4s in preview-only trial (128 trials) or was replaced by a search scene (400 ms, remaining 128 trials). For the search trials, a response screen then prompted participants to indicate whether the target was present or absent by using a button box with their right hand (max response duration 1.5 s). Feedback was provided after every search trial by a red or green dot at fixation. After every run, participants were additionally given feedback on their average performance.

Target category and scene context changed on a trial-by-trial basis and the factors Trial type (search, preview only), Target presence, Target category (Book, Bowl), Scene context (blue, yellow living room) and Position of the associated anchor (left/right) were all counterbalanced within each run.

### Localizer task

To localize voxels that were selective to the targets and/or anchors, we also included four runs of a separate localizer task. In this task, participants saw the anchor and target objects used in the main task, now presented in isolation on a grey background. There were either two identical objects shown in the periphery (at the location of the table centers in the search task) or a single object in the center of fixation (as they would appear after an eye movement to fixate them). Each run consisted of 32 miniblocks (4 repetitions of 8 conditions: book/bowl/Table 1/Table 2 x central/ peripheral location).

Within each miniblock, participants saw 8 stimuli of the same category (0.4s on, 0.6 s off, 2s fixation baseline after each miniblock) and had to respond to size-oddball targets, in which the stimuli appeared slightly larger than usual (i.e., deviating from their typical, memorized size; 1.15x size increase for anchor tables, 1.5x for target objects) by pressing a button. There was one such oddball target per miniblock and participants received feedback about the percentage of found targets after each block. For target miniblocks, different exemplars were shown within a miniblock. As this was not possible for the anchor tables, we introduced slight rotations along their vertical axes in the images, to avoid a pixel-based strategy for the task and decrease repetition suppression effects.

The localizer task was presented interleaved with the search task, with one localizer followed by two search task runs. For the first three subjects, a different localizer task was used. While they saw the same stimuli (without oddball size targets), they were asked to respond to a brief rotation of the fixation cross. All other presentation parameters were the same for all participants.

Of note, for all analyses, we did not observe generalization (i.e., successful cross-decoding) between visually evoked responses in this localizer and the preview trials of the main task. This may be due to insufficient statistical power to detect such a generalization or that different features were relevant to discriminate objects in the periphery for quick guidance of eye movements and comparing objects to their memorized size in this localizer task, resulting in less overlap across tasks (Henderson et al., 2023; McKee et al., 2014). Another possibility is that preparatory activity in our task was overall less sensory-like (Gong et al., 2022). As such, this null-finding is difficult to interpret, and our analyses focus on cross-validation across runs of the main experiment and cross-decoding across different scenes, agnostic to the representational format of the preparatory template.

### Anchor-target association training

Immediately prior to scanning, participants practiced the search task and learned to associate anchors and targets. They saw example target objects and search scenes and practiced the search task (32 trials). During this practice they were informed that within each of the two rooms, each target was associated with one of the two tables in the room, and that it would always appear on the respective table if it was present.

After this practice, participants were tested on the associations using a 2AFC-task (12 trials). On each trial, they were shown one of the living rooms with empty tables left and right and the name of a target written in the image center, and had to indicate the location of the associated table given target and scene-context using a keyboard. Performance on this task was 83.09% (sd 14.72). Participants received feedback after each trial and were reminded of the associations after the task if needed.

### Stimuli

640 3D rendered search scenes were created using Blender 2.92 (Blender Foundation) and the Eevee rendering engine. These scenes depicted a blue or yellow living room with identical spatial dimensions and camera positions, but different furniture.

In both rooms, two visually distinct tables were placed in the foreground, slightly below fixation, on the left and right. Four objects were placed on each table, each being either one of the target objects (books and bowls, drawn from 20 unique exemplars per target category, varying in color and size), or non-target objects that could be plausibly found on living room tables (40 objects, e.g., a tea mug, chess board, baseball cap, headphones, game controller, vase, candles, laptop). 320 unique object constellations were created - 20 exemplars for each of combination of counterbalancing factors (scene context, target presence, associated anchor left or right) and the associated anchors (counterbalanced across participants) added. Models for furniture, targets, and distractor objects were taken from https://sketchfab.com/or newly built/modified. Custom python scripts ensured objects did not intersect with each other, and images were manually checked to ensure objects did not significantly occlude each other. Search scenes subtended about 25 x 12.62° (20 x 11.25° for the first three participants).

Scene images with blurred tables (both tables averaged and pixelated left and right) were used as masks after the search scene, and two scenes with grey occluder squares in front of the tables used as preview.

For the localizer, isolated target and anchor objects were rendered without the scene background, either at the peripheral locations the anchors occupied in the search scene or in the center at the same height. For the target objects, 20 images per localizer condition showed all category exemplars used in the search task. For the anchor objects, 10 images were created per condition, showing the same table slightly differing in its rotation along the vertical axis (equidistant steps from -10 to 10 degrees from their typical orientation in the search scenes).

### Setup & Eye tracking

Inside the scanner, stimuli were presented on a 32 inch IPS BOLD screen (1920 x 1080 pixels, 120 hz refreshrate) placed at the back of the scanner bore, and could be viewed by participants through a mirror mounted on the head coil.

During scanning, participant’s left eye was tracked by an Eyelink 1000+ eye tracker (SR Research, sampling rate 1000 Hz). At the start of the experiment, the eyetracker was calibrated using a 9-point calibration procedure (resorting to a 5 point calibration in case no accurate 9-point calibration was possible) and recalibrated if required.

### Eye tracking AOIs

Three rectangular areas of interest (AOIs) were defined for analyzing fixations during the search scene. First, a center AOI around the fixation cross (4.5 x 10.5°) to ensure fixation was in the image center before the search scene appeared and one centered on the left and right anchor table each (7.5 x 10.5°). On 94.15% (sd 9.65) of trials, at least one fixation was recorded during presentation of the search scene, and 27.08% (sd 10.32) of trials had two or more recorded fixations. 83.33% (sd 19.36) of trials with a fixation also had an initial fixation in the center AOI at scene onset, of which 80.52% (sd 10.76) of first fixations were then directed at one of the anchor AOIs.

### fMRI acquisition and preprocessing

Data were acquired on a 3T Siemens SKYRA Scanner using a 32-channel head coil. A T2-weighted gradient echo EPI sequence was used for acquisition of functional data (TR 1.5 s, TE 33.4 ms, flip angle 75°, 2 mm isotropic voxels, 68 slices, 4x multiband acceleration factor). For the search task, 198 images were acquired per run and 235 images were acquired per run for the localizer runs. A high-resolution T1-weighted anatomical scan was acquired prior to the experimental runs, using an MPRAGE sequence (TR 2.3 s, TE 3.03 ms, flip angle: 8°, 1 mm isotropic voxels, 192 sagittal slices, FOV 256 mm).

Data preprocessing was performed using SPM12 (https://www.fil.ion.ucl.ac.uk/spm/). Preprocessing steps included spatial realignment, co-registration of functional and anatomical scans and normalization to MNI 152 space. A Gaussian filter (FWHM 3 mm) was then applied to smooth the images.

Subject level GLMs were estimated on the preprocessed images. For the search task, the 4 combinations of targets and scene context were modelled as regressors of interest, by convolving a boxcar function spanning from preview onset to offset with the canonical HRF curve provided in SPM12. Importantly, this model only included preview-only trials.

For the localizer runs, individual miniblocks were modelled as boxcar functions spanning the duration of a miniblock and were also convolved with the canonical HRF curve. There were 8 conditions of interest: books, bowl, anchor table 1 and anchor table 2 each presented centrally or peripherally.

Six motion regressors and one run-based regressor were included as nuisance regressors and all betas were estimated on a run-based basis.

### ROI definition

Two main ROIs (EVC, LOC) were defined for each participant. ROI definition was based on group level masks (EVC: Brodmann areas 17, 18, LOC: taken from (2012)), resliced into MNI space. To exclude voxels of non-interest (e.g., white matter voxels, visually unresponsive voxels), these group-level masks were intersected with contrasts from the localizer task for each participant, as follows: within each ROI, we selected voxels that discriminated between the targets or anchor objects, when they were visually presented during the independent localizer task, by intersecting the group levels masks with respective F-contrasts (book vs. bowl or table 1 vs. table 2). This was done separately for each hemisphere, to be sensitive to potential hemispheric differences. For each contrast, hemisphere and brain area, we first created one large ROI mask (unrestricted -ROI) including all active voxels based on that contrast (p < 0.05, uncorrected) by intersecting the respective group-level masks and participant contrast maps. In a last step, we combined the masks created for each contrast, thus including both anchor and target-discriminating voxels within one ROI-mask, and results are based on these combined ROI maps. Averaged across participants and hemispheres, these masks included 1289.85 voxels in EVC (sd 58.17) and 917.32 voxels (sd 50.72) in LOC. This procedure ensured we included informative voxels, in which any preparatory bias for targets or anchors should be strongest, while still basing our analyses on a large enough subset of voxels.

To ensure that our results were robust to different ROI definitions, we also created smaller sub-ROIs for each brain area and participant. Five sub-ROIs were created, including an equal number of the top x most target and anchor selective voxels within each hemisphere in equidistant steps up to 424 voxels for EVC and 264 voxels per hemisphere for LOC. Those numbers reflected the median number of significantly target-selective voxels across hemispheres and participants (always lower than the number of anchor-selective voxels). All our main results were also consistent when considering the target- or anchor-selective voxel ROIs in isolation (see Figure S1).

### Multivariate pattern analysi*s*

All multivariate analyses were performed using linear support vector machines (SVMs), using The Decoding Toolbox (TDT: Hebart et al., 2015) and the libsvm library. Classifiers were trained and tested on the run-based beta-weights after GLM estimation. For every ROI and voxel count, separate classifiers were trained for each hemisphere. As we did not observe any hemispheric differences, the final classification performance was then averaged across hemispheres before statistical testing (see Figure S2 for decoding results within individual hemispheres).

Classification was either performed in (1) a leave-one-run out 8-fold cross-validation scheme to test for target or target-associated anchor information within the search task runs (training the classifier to, e.g., distinguish all book and bowl trials), or (2) a cross-classification scheme to test for generalization across scene contexts (training on search task preview trials of one scene context and testing on the other). For cross-classification, the classification directions were averaged before statistical testing.

### Searchlight analysis

To complement the previous ROI analyses, we ran an additional searchlight analysis, testing for generalization of the anchor template across scenes (1-sided test). Similar to the ROI analyses, we first identified target- and/or anchor-selective voxels by running additional searchlights (restricted to a mask of cerebral cortex regions based on the AAL atlas (2002)) to decode the targets or anchors from the localizer betas. Voxels with significant decoding for either the target or anchor at the group-level (alpha = 0.05) were then used to create a mask for the cross-decoding searchlight. For all searchlights, spheres had a radius of 5 voxels (resulting in around 524 voxels in total).

### Statistical tests

To test for differences between conditions in the behavioral/eye tracking analysis or to determine above-chance decoding in the all-ROIs, we performed bootstrap tests against chance, resampling individual participants with replacement for 10000 iterations. P-values reflect 2-tailed tests unless indicated otherwise.

Threshold free cluster enhancement (TFCE) was applied to the sub-ROI data and searchlight analyses, using the CoSMo MVPA toolbox (Oosterhof et al., 2016). This procedure boosts belief in neighboring data points containing signal, and was applied both to real as well as synthetic null-data. The final p or z-values reflect how likely a given TFCE score is, given the maximum TFCE scores across the null-data and thereby accounts for multiple comparisons.

## Acknowledgments

We thank Cara Struckmann for help with stimulus creation, Giacomo Aldegheri for help with Blender programming, and Surya Gayet for helpful comments on this manuscript.

## Author Contributions

Conceptualization: ML, FPL, MVP

Methodology: ML, FPL, MVP

Software: ML

Formal Analysis: ML

Investigation: ML

Resources: ML

Data Curation: ML

Writing—original draft: ML, FPL, MVP

Writing—review & editing: ML, FPL, MVP

Visualization: ML

Supervision: FPL, MVP

Funding Acquisition: MVP

## Data and materials availability

Behavioral, eyetracking and anonymized fMRI data, stimuli and experiment/analysis code can be accessed on the Radboud Data Repository (https://doi.org/10.34973/dg68-4p18).

## Supplementary Materials

### Behavioral analyses

Overall behavioral accuracy in the search task was 72.46% (sd 8.91). Hit rate was 66.16% (sd 12.61), with 21.16% (sd 10.89) false alarms. Sensitivity (*d’*) was 1.51 (sd 0.61) and criterion (*c*) 0.21 (sd 0.24) on average.

Sensitivity (*d’*) was 1.51 (sd 0.61) and criterion (*c*) 0.21 (sd 0.24) on average. Thus, as intended, finding the targets in the search task was challenging, but not impossible.

We established whether performance was matched for both targets and scene contexts, using a Target (Book, Bowl) x Scene Context (yellow, blue living room) ANOVA. Participants were equally accurate for both targets and both rooms (no main effect of Target: F(1,33) = 1.04, p = 0.32, η_p_^2^ = 0.03; no main effect of Scene context: F(1,33) = 0.07, p = 0.79, η_p_^2^ < 0.01; no Target x Scene context interaction: F(1,33) = 3.06, p = 0.07, η_p_^2^ = 0.09). Next, we tested whether performance differed for search on the two anchor tables, using an Anchor (Table 1, Table 2) x Scene Context (yellow, blue living room) ANOVA. Participants were more accurate on the smaller and brighter table (F(1,33) = 12.85, p < 0.01, η_p_^2^ = 0.28), possibly due to the better contrast between the table and the objects placed on it. However, this advantage depended on the context, as indicated by a significant Anchor x Room interaction (F(1,33) = 5.84, p = 0.02, η_p_^2^ = 0.15). While performance for the two tables differed in the yellow living room (CI = [4.86, 10.85] p < 0.001) they did not differ for the blue living room (CI = [-3.2, 5.19], p = 0.58). Altogether, these findings indicate that there were no consistent accuracy differences across targets, anchors, or rooms.

**Fig. S1.**
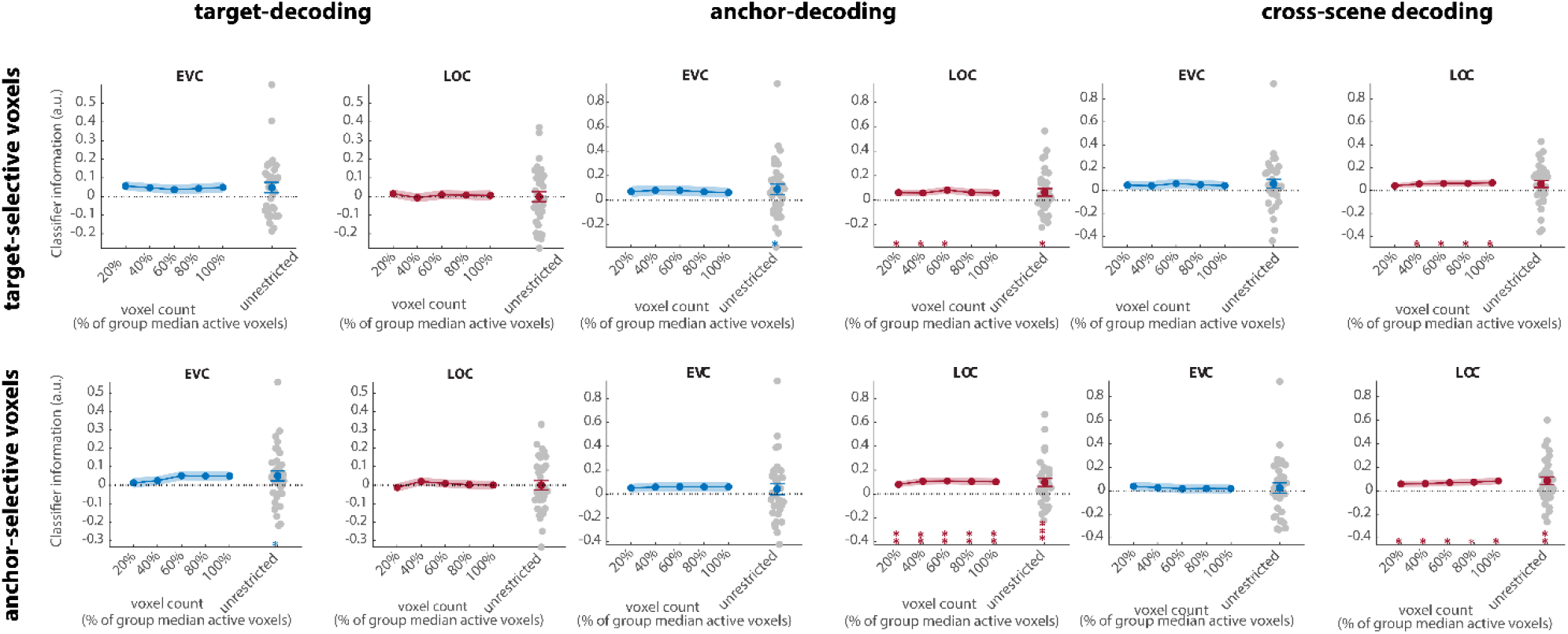
Decoding results shown separately for target and anchor-selective voxels. Decoding in EVC and LOC for all selective voxels of individual participants (unrestricted-ROI) and sub-ROIs of different sizes. Grey dots show decoding for individual participants in the largest (unrestricted) ROI. All error bars are SEM. *, p<0.05; **, p<0.01

**Fig. S2.**
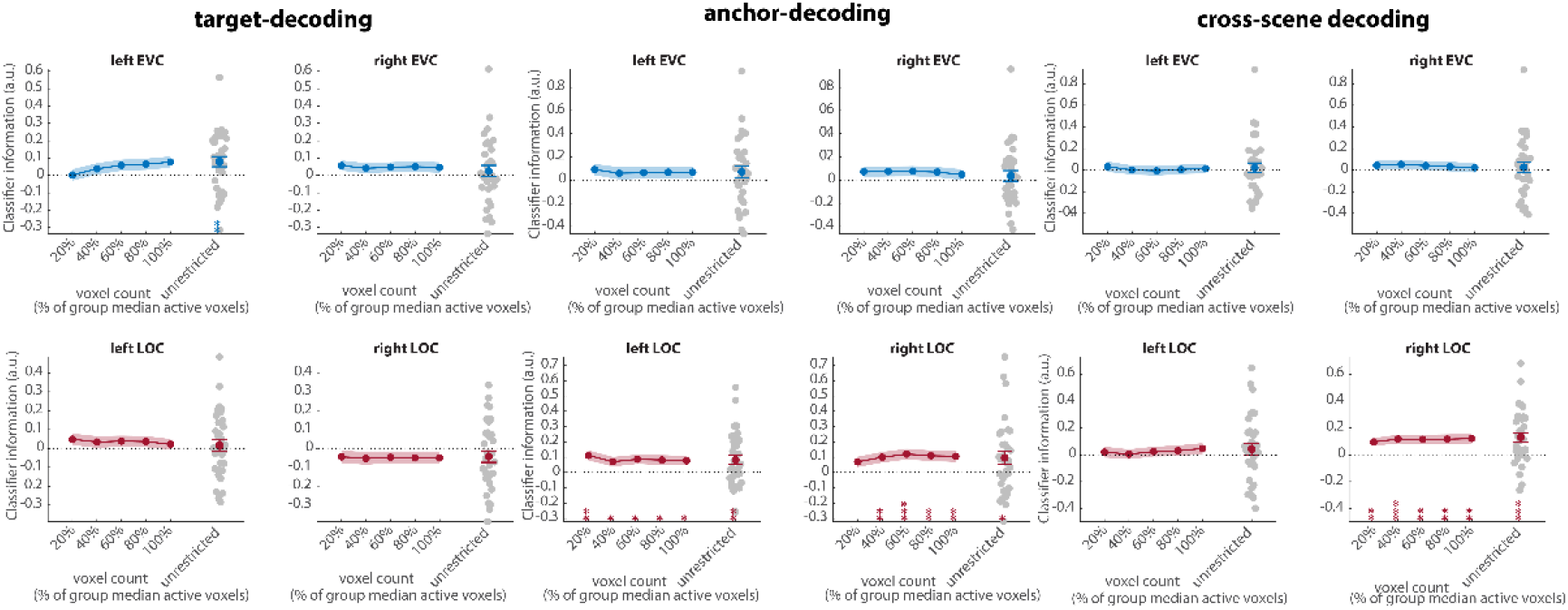
Decoding results shown separately for left and right hemisphere. Decoding in EVC and LOC, for all selective voxels of individual participants (unrestricted-ROI), and sub-ROIs of different sizes. Grey dots show decoding for individual participants in the largest (all) ROI. All error bars are SEM. *, p<0.05; **, p<0.01;***, p<0.001

**Fig. S3.**
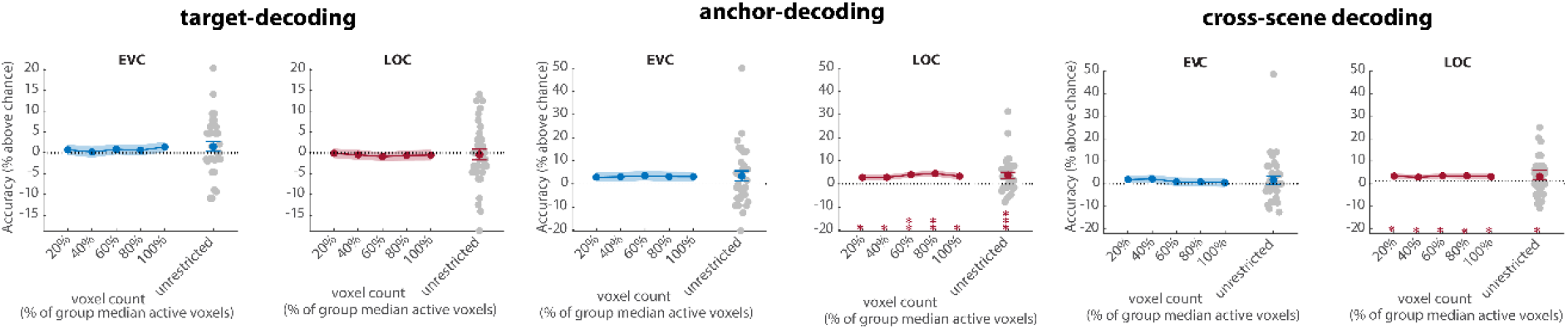
Decoding results using decoding accuracy. Decoding in EVC and LOC for all selective voxels of individual participants (unrestricted-ROI), and sub-ROIs of different sizes. Grey dots show decoding for individual participants in the largest (all) ROI. All error bars are SEM. *, p<0.05; **, p<0.01; **, p<0.001

**Fig. S4:**
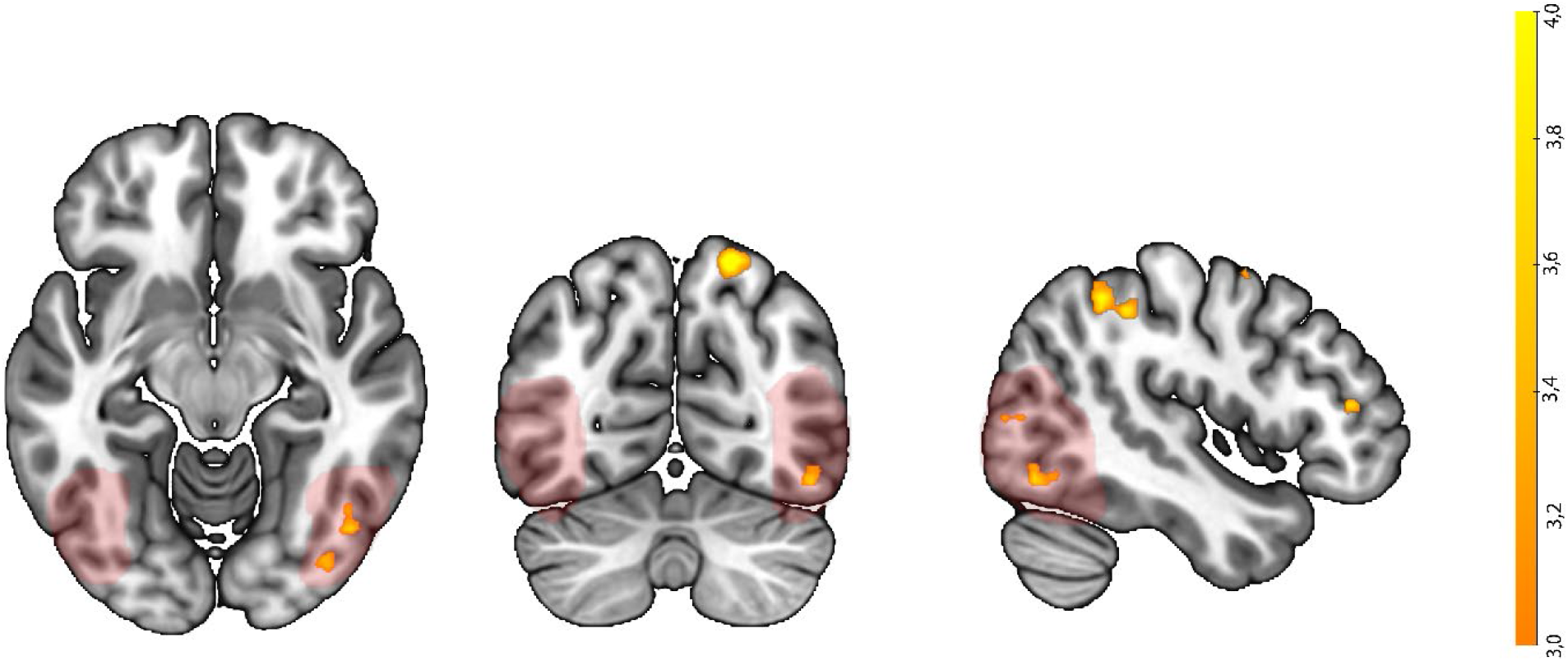
Uncorrected whole-brain searchlight (no voxel selection) for decoding the target-associated anchor across scene-contexts. The searchlight shows voxel clusters within the LOC mask (marked in red). Searchlight is thresholded at p = 0.005 (2-sided). Color indicates t-values without threshold-free cluster enhancement (TFCE).

